# Controlling T cells shape, mechanics and activation by micropatterning

**DOI:** 10.1101/2020.09.15.295964

**Authors:** A. Sadoun, M. Biarnes-Pelicot, L. Ghesquiere-Dierickx, A. Wu, O. Théodoly, L. Limozin, Y. Hamon, P.-H. Puech

## Abstract

We designed a strategy, based on a careful examination of the activation capabilities of proteins and antibodies used as substrates for adhering T cells, coupled to protein microstamping. This allowed us to control at the same time the position, shape, mechanics and activation state of T cells. Once adhered on shaped patterns we examined the capacities of T cells to be activated with soluble aCD3, in comparison to T cells adhered to a continuously decorated substrate with the same density of ligands. We show that, in our hand, adhering onto an anti CD45 (aCD45) antibody decorated surface is not affecting T cell calcium fluxes, even adhered on variable size micro-patterns. We further demonstrate this by expressing MEGF10 as a non immune adhesion receptor in T cells to obtain the very same spreading area on PLL substrates and Young modulus than immobilized cells on aCD45, while retaining similar activation capabilities using soluble aCD3 or through model APC contacts. We propose that our system is a way to test activation or anergy of T cells with defined adhesion and mechanical characteristics, and may allow to dissect fine details of these mechanisms since it allows to observe homogenised populations in standardized T cell activation assays.

## Introduction

The engagement by the T lymphocyte receptor *αβ* (TCR, T cell receptor) of molecules of the major histocompatibility complex loaded with agonist peptide (pMHC) is a central step in the adaptive immune response ^1^. This recognition is characterized by its high specificity and sensitivity ^2^, with a broad spectrum of intracellular signaling events leading to T cell activation and effector functions ^3^. How T cells contrasting cellular responses like clonal expansion, anergy or apoptosis reflect the quantitative and qualitative diversity of antigens remains largely an enigma from a molecular and kinetic point of view ^4–7^.

However, the main protein protagonists have been identified and extensively documented on both the antigen presenting cell (APC) and the T cell. At a minimum ^8^, TCR*αβ* is an octameric polypeptide complex, composed of a clonotypic and hypervariable recognition module (chains α, *β*) and a transduction module formed by the CD3 invariant chains associated in dimers ^9^. The binding of TCR to pMHC triggers the phosphorylation of ITAM units, located in the cytoplasmic domains of the CD3 subunits, by the kinase of the Src Lck family. This results in the recruitment and activation of kinases from the ZAP-70 Syk family, which in turn phosphorylate various critical signaling intermediates, including the LAT adapter ^9,10^. A major gap in this picture is the lack of understanding of the mechanism by which the engagement of the TCR/CD3 ligand on the T cell surface causes the phosphorylation of CD3 chains by the Lck kinase located on the inner sheet of the plasma membrane, a process that is commonly referred as “TCR triggering”. Various models have been proposed so far to explain this ^11^, including the molecular aggregation, pseudo-dimer, kinetic segregation or conformational change models ^2,12,13^.

While all these different models are not fully consistent with each other, some of them may represent different facets of the same mechanism that are simply shifted in time or space. Finally, all these models attempt to explain the profound paradox that exists between, on one hand, the selectivity / sensitivity / rapidity of responses contrasting, on the other hand, with the diversity / scarcity of antigens exhibiting low affinities (as typically obtained by “3D vs. 2D” surface plasmon resonance) for TCR ^4,14^.

Seminal studies ^15–17^ measuring the kinetic parameters of 2D TCR/pMHC interactions (in a membrane environment) radically contradict studies conducted with molecules in solution and demonstrating very rapid association/dissociation cycles of the same TCR for the same pMHC, the affinity of TCR for its natural ligand being of the same order as the one between adhesive molecules such as ICAM1 and LFA1 ^18^.

All these parameters classify TCR at the heart of a paradigm of unconventional interactions between membrane receptors / ligands. Ultimately, the transverse organization of the plasma membrane (its composition, its inhomogeneities, the nature of its interactions) would influence the ability of T lymphocytes to respond to the antigen ^19^.

Recently, and driven by technological, biophysical and biological developments, forces that may be at play at the interface of the T cell and APC while the former scans the latter have been proposed to be crucial for T cell recognition and subsequent activation ^18–21^. Experiments at single molecule scale have tested if TCR/pMHC bond rupture forces ^22^ or lifetimes ^23,24^ depends on the quality of the peptide and have proposed that this bond could exhibit a behaviour more complex than expected, namely a catch bond behaviour ^23^, where, depending on the peptide, the lifetime of the bond can be modulated and even extended for a range of forces compatible with the ones that cell protrusions may exert ^25,26^. The geometry of the force application has also been investigated using refined micromanipulation methods ^27–29^. Moreover, the way the forces are exerted, continuously vs. intermittently, have been proposed to be an important modulator of recognition ^23,30^, directly involving the T cell cytoskeleton in the recognition ^31^. Aside, at single cell level, T cells have been observed to react differently as a function of the mechanics of the substrate they are interacting with ^32,33^, in line with the recent demonstration that APCs exhibit different mechanical properties, which are modulated along the inflammation process ^34^. The complexity of the response to substrate mechanics, modulated by the molecules used to make the cells spread on it, have been very recently examined ^35^. Aside from cell mechanics, cell shape has been shown to have a non negligible impact on cellular functions ^36^ and T cell activation ^37^. Applied forces can also, in turn, affect the shape of the T cell itself ^38^ and APC cytoskeleton may impact T cell recognition ^39–41^.

Single cell fluorescence studies ^42^ or force based measurements where cells are adhered on a substrate ^22,30,31,37,43^ or using single T cell / APC interaction ^44–46^, while highly informative, have the inherent drawback of relying on the parallel or more often sequential measurement of cells in a heterogeneous population, which may possess slightly different mechanics and shape. In this frame, we recently proposed a technique to couple force based measurements using atomic force microscopy and optical tracking of calcium fluxes or opto-genetics perturbation of cell mechanics as a new tool ^43^.

Here, we designed a strategy to “standardize T cells”, based on a careful examination of the activation capabilities of proteins and antibodies used for adhering T cells on micro-stamped substrates ^47^. We applied it on a T cell line recapitulating activation steps up to Interleukin 2 production, independently of CD28 costimulation ^42^. It allowed us to control at the same time the position, shape, mechanics (as measured using atomic force microscopy indentation) and activation state of T cells (as followed by calcium fluxes). We examined the capacities of activation, using soluble aCD3, in comparison to T cells adhered to a continuously decorated substrate with the same density of ligands, and verified that T cells have a similar behaviour in regard to classical activation assays. We further show that, with our system, CD45 mobilization on continuous substrates or patterns of controlled areas is not affecting the proper activation as observed through calcium fluxes. To confirm this, we expressed an exogenous adhesion molecule in T cells that can be used for immobilisation purpose and demonstrated that this did not perturb T cell calcium response similarly as our anti CD45 strategy, hence suggesting this new construct as a new tool to dissect T cell activation for biophysical techniques which require adherent cells.

## Material and methods

### Chemicals

Chemicals and proteins were obtained from Sigma Aldrich except when noted below. Culture media and supplements were obtained from Gibco. References can be found in ^42,43^ for generic components.

### Cell culture

#### Culture

3A9m T cells were obtained from D. Vignali ^48^ and cultured in RPMI completed with 5 % Foetal Bovine Serum (FBS), 10 mM Hepes in 5 % CO2 atmosphere. COS-7 APC were generated as previously described ^42^ by stably co-expressing the a and the *β* chains of the mouse MHC class II I-Ak, cultured in DMEM (5 % FBS, 1 mM Sodium Pyruvate, 10 mM Hepes, and geneticin 10µg/ml). Cells were trypsinized up to three times a week by treating them with either Trypsin/EDTA or 0.53mM EDTA at 37°C for up to 5 min.

#### Cell fixation & labelling for confocal imaging

3A9m cells were fixed using 4% Paraformaldehyde (20 min on ice), saturated with glycine 0.1M and permeabilized with 0.1% Triton and labelled in presence or not of 0.01mM Hoechst (at room temperature, 10 min) with 0.5µM phalloidin-Alexa Fluor 488 (Life technologies) (at room temperature, 20 min).

#### Protein expression

The expression levels for TCR and CD45 (for T cell hybridomas), and MHC (for COS-7 APC) were routinely monitored by flow cytometry (BD Biosciences, LSR2) once a week. Expression of CD4 was tested once for every cell batch since it was observed to be stable over time (not shown).

### Antibody production and labelling

#### Production

The antibodies used for this study were produced from the hybridoma collection of Centre d’Immunologie de Marseille-Luminy (CIML) (namely anti TCR H57.597, anti CD3 2C11, anti CD4 GK1.5, anti CD45 H193.16.3, anti MHC II 10.3.6). Briefly, hybridoma were routinely grown in a complete culture medium (DMEM, 10% FBS, 1 mM sodium pyruvate) prior to switching to the expansion and production phase. They were cultured in DMEM with decreasing concentrations of low immunoglobulin FBS (Life Technologies) down to 0.5%. Cells were then maintained in culture for 5 additional days enabling immunoglobulin secretion prior to supernatant collection and antibody purification according to standard procedures.

#### Fluorescent labelling, when needed

200µg of antibody solutions (in PBS 1X) were mixed with N-Hydroxysuccinimide (NHS) ester functionalized dye (dye/protein ratio set to 3:1) with sodium bicarbonate solution 0.1M (pH 8.3). We used Alexa Fluor 488 and 647 (Thermofisher Scientific) or Atto 565 (Sigma Aldrich). After incubation for 1h at RT under constant agitation, uncoupled dyes were separated from labelled antibodies by size exclusion chromatography (on Sephadex G25 PD10 columns GE Healthcare) followed by dialysis against PBS 1x overnight at 4°C. At last, a dialysis against PBS 1x was performed for 1hr. The concentration and labelling efficiency was then assessed on a Nanodrop 100 (Thermo Scientific). The obtained dye/protein ratio was typically between 1 and 2.

### MEGF10 stably expressing T cells

The MEGF10::EYFP construct has been previously described in ^49^. Briefly the 220 last bases of the 5’ UTR together with the human cDNA coding for MEGF10 were subcloned upstream and in frame with the EYFP sequence into the pEYFP-N1 vector. The plasmid was introduced into 3A9m T cells by AMAXA nucleofection (Kit V, B024; Lonza) according to the manufacturer and as already detailed in ref. ^50^.

### Continuous substrates

#### LabTek chambers for calcium imaging

8 well-LabTek II chambers (Nunc) were coated by adsorption of solutions of antibodies in PBS 1x overnight at 4°C, then saturated with PBS 1X, 1% BSA solution, for 1h at 37°C prior to observations.

#### Glass bottom Petri dishes

##### Antibodies Coating

Glass bottom Petri dishes (WPI Instruments, Fluorodish FD35-100) were incubated with 50µL of aCD45 H193.16.3 (final 50µg/ml) for 45 minutes at room temperature. Once the droplet was removed, the surfaces were extensively rinsed first with sterile PBS 1X, then with HBSS-H.

##### Poly-L Lysine coating

The petri dishes were activated with residual air plasma for 10 minutes, then incubated with a mix 9:1 poly-L-lysine:Alexa Fluor 546-labelled poly-L-lysine solution (0.1% or 0.01% w/v H_2_O, Sigma P8920-100ml, labelled with Alexa Fluor 546 NHS Ester Life Technologies A20102 following provider protocols) for 45 minutes at room temperature, and rinsed as above.

### Protein microstamping

#### Stamps production

Stamps were produced using soft photolithography techniques from in-house designed patterns. They consist of regularly spaced disks of size between 5 and 50µm in diameter. The spacing was optimized so that one T cell could not adhere to more than one pattern. For the smaller sizes, immobilization was not very efficient for us (eg. 5µm), while for larger ones (eg. 50µm), several cells may adhere to the same pattern (not shown).

To produce the stamps, we used PDMS (Sylgard 184, Dow Corning) in a ratio w/w 1:10 of curing agent. 1cm^2^ stamps of typically 0.5cm thickness were then cut using a scalpel. The stamps were then washed in two steps in a sonicator at max power : (i) 20min at 65°C with 5 % Decon; (ii) 20min at 65°C in ultrapure water. Finally, they were dried under gentle nitrogen flux and stored protected from dust before immediate use.

#### Stamping procedure

50µL of the desired stamping solution (typically 10µg/mL – 100µg/mL of protein) was spread onto the stamp and let adsorb during 45 min at room temperature, protected from light. The droplet was then aspirated and the stamp briefly dried using a nitrogen flux, at room temperature. The stamp was immediately gently pressed using tweezers on a clean glass bottom Petri dish and the transfer was let to occur for 10 min in a cell culture incubator at 37°C, protected from light. After removing the stamp with great care, the substrate was intensively rinsed with 0.2 µm filtered PBS 1x before adsorption of the desired complementing molecule (typically 10µg/mL – 100µg/mL of protein, with an optimal eg. for aCD45 at 50µg/mL when the first one was (b)BSA, see text) for 1hr at room temperature, protected from light.

Final extensive rinsing is performed with the desired experimental buffer (here, HBSS/1 mM Hepes) and the final volume in the Petri dish was set to 2-3 mL for further experiments. Whenever possible, the quality of the stamping was checked by fluorescence either of the stamped molecule or of the complementary one, or after adding eg. fluorescent streptavidin over biotinylated substrates.

#### Cell adhesion on stamps

Cells were seeded over the stamped region and let to adhere and spread for 1 hour in a cell culture incubator, protected from light. After observation in transmission, fluorescence and eventually interference microscopy (RICM, see below), the non adherent cells were removed by a gentle rinsing step with HBSS and the remaining adhered cells were let to rest for at least 15 minutes before starting any experiment.

### Calcium imaging

The measurements of CD4+ T cell cytosolic calcium fluxes were performed according to the “methods for automated and accurate analysis of cell signals” (MAAACS) extensively described in ref. ^42,50^.

#### Calcium reporter loading

In brief, T cells were loaded with PBX calcium reporter (BD) diluted in 1X dye loading solution at 37°C for 1 hour in the dark, according to manufacturer’s instructions. Cells were then washed twice by gentle centrifugation and resuspended in Hank’s balanced salt solution buffered with Hepes (1mM) (HBSS-H). They were then introduced in the desired observation chamber (Labtek or glass bottom Petri dish).

#### Confocal microscopy

Analysis was performed using a Zeiss LSM 780 confocal microscope equipped with a C-Apochromat 40X/1.2 water immersion objective as well as an argon laser with a 488 nm dichroic and a 505-530 nm band pass filter. A temperature control system was used to ensure that 37°C was maintained during the entire acquisition. Time-lapse movies were typically made of 150 images taken every 7 seconds (with a pinhole set to 4 airy units), while cells were kept at 37°C using a hot plate. When needed, a similar second serie of acquisitions was launched upon activation with soluble anti CD3 antibodies (2C11; 20µg/ml) or a suspension of COS APC cells loaded or not with the desired peptide (1 million cells/ml).

#### Quantification

Calcium response intensity parameters as well as velocity and cell shape (as “shape index”) were determined by MAAACS as previously described ^42^. Specific calcium response amplitude was determined upon specific thresholds (depending on experimental stimuli) set according to our previous studies. The percentage of responding cells, average response amplitude and average response time fraction were automatically calculated and tabulated in MS Excel data sheets. Results were plotted as dot plots, limited to calcium responses above threshold.

### Reflexion interference contrast microscopy (RICM)

A Zeiss Axiovert, equipped for fluorescence microscopy (GFP, TRITC filter sets) and a specific Antiflex 63x objective (Zeiss) was used to monitor the contact zones of cells with substrates ^51–53^. The RICM was performed at 546 nm wavelength by using a dedicated filter set. Dark zones in the images were the signature of close apposition of the membrane, to the few nanometers scale, to the substrate and quantified using either hand selection or homemade macros in Fiji/ImageJ software ^54^.

### AFM set-up

The set-up has been described in details elsewhere ^43^. Measurements were conducted with an AFM (Nanowizard I, JPK Instruments, Berlin) mounted on an inverted microscope (Zeiss Axiovert 200). The AFM head is equipped with a 15 µm z-range linearised piezoelectric scanner and an infrared laser. The set-up sits on an active damping table (Halcyonics). AFM was used in closed loop, constant height feedback mode. Temperature control was achieved using a Petri Dish Heater module (JPK Instruments), which controller was connected to the AFM one.

Bruker MLCT-UC cantilevers were used in this study, and glass beads (5 µm in diameter, silica beads from Kisker Biotech GmbH, larger than cantilever tip) were glued at their extremity using micropipette micromanipulation with UV optical glue (OP-29, Dymax) and intense UV curing (10 min at maximal power of a UV oven (BioForce Nanosciences)). The sensitivity of the optical lever system was calibrated on the glass substrate and the cantilever spring constant by using the thermal noise method ^55^, using JPK SPM software routines (JPK Instruments, version > 3) *in situ* at the start of each experiment. The calibration procedure for each cantilever was repeated three times to rule out possible errors and spring constants were found to be consistently close to the manufacturer’s nominal values and the calibration was stable over the experiment duration.

The inverted microscope was equipped with 10x, 20xNA0.8 and 40xNA0.75 lenses and a CoolSnap HQ2 camera (Photometrics). Bright field images were used to select cells and monitor their morphology during force measurements. For fluorescence, the microscope was equipped with a LED illumination system (Colibri 2, Zeiss) and suitable filter sets. Images were obtained through either Zen software (Zeiss) or µManager ^56^.

### T cell mechanics using AFM

In order to measure the Young modulus of the shaped and non shaped cells, AFM indentation experiments were performed using MLCT-UC levers on which a bead, smaller than the cell, was glued (see above). Cells were adhered at 25°C when loaded with calcium reporter dye to avoid compartmentalization and 37°C overwise ^50^ and were selected by bright field examination and, if needed, the protein printed pattern was examined using fluorescence microscopy. The indentation experiments were then performed at a prescribed, constant, temperature (25 or 37°C). The occurrence of dye compartmentalization ^57^ limits the duration of the experiment to 1h30.

To start, the bead was positioned as much as possible above the centre of the cell. The maximal force to be applied was set at 500pN (leading to indentation depths of the order of one µm, smaller than the typical cell size), the contact duration from zero (pure elastic deformation, the most common experiments) to up to 10 sec (to follow the viscous relaxation), the speed of pressing and pulling at 2µm/s in all cases for a 7µm displacement. Depending on the measurement, a single or up to 10 force curves were recorded for each cell, with a delay time of 1 sec for repeated acquisitions in order to let the cell recover. Data was recorded at 2048 Hz.

For determination of the Young modulus, each force curve was examined by eye and processed with the “Hertz model procedure” included in JPK DP software (JPK Instruments)^58^ : corrections for baseline, possible tilt of the baseline and a classical Hertz model for spherical tips were applied, making the hypothesis that the cell behaves as an incompressible material ^59^. Only a subset of the entire force span (from the baseline to the maximal contact force) was fitted : we chose to fit over 0.5µm of indentation to minimize contributions from the nucleus (see Results). Young modulus were found to be coherent with published ones for T and immune cells ^34^. For the evaluation of the force relaxation, we estimated a power-law exponent *n* for *f(t)* as *f=f*_*0*_ *(t/t*_*0*_)^*n*^ following ^60^. We verified that, when maintaining the piezo position constant, the relaxation induced only a moderate (∼ 4-5 % in average) variation of the cell’s indentation. Then, we applied an *ad hoc* fit (using a dedicated Python procedure) and extracted *n*.

For each parameter, a median value per cell was then calculated and tabulated in each condition. The resulting box-plots were then plotted using Python libraries Matplotlib (matplotlib.org/) and / or Seaborn (stanford.edu/∼mwaskom/software/seaborn/). We validated this way of pooling the data experimentally since no obvious correlation between the Young modulus and the force curve number (corresponding to the « mechanical history » of the cell) was observed (not shown).

### Data processing and statistics

Data is presented as mean +/- SEM overwise precised. Data plotting and significance testing were performed on Linux or Windows PC 64 bits machines using Python (packages Seaborn, Matplotlib, Scipy, Numpy, Scikit) and/or R / Rstudio (http://cran.rstudio.com/, packages asbio and pgirmess for Kruskal-Wallis or Wilcoxon tests) and/or Graphpad Prism (6 or 7).

## Results and discussion

The build-up of a system where T cell localization, shape, mechanical properties and activation level were controlled was divided into successive steps : (i) identifying relevant molecules to immobilize cells on a surface with cells keeping a low induced activation while remaining activable eg. using soluble molecules, (ii) determining a reproducible micropatterning protocole for shaping small and reactive T cells in the frame of (i), (iii) characterizing their mechanical properties such as elasticity, (iv) comparing their activability to more classical conditions and (v) qualifying novel methods of immobilization using a new molecular construct that can be easily transfected into T cells.

### Optimizing T cell adhesion while controlling their activation level

T cells are mostly non-adherent cells in culture, so that the first step was to have them adhere on glass substrates, considering that the substrate should immobilize the T cells, shape them while keeping their biological properties to be activated. Adherent and repellent molecules were tested, in order to create afterwards adhesive patterns surrounded by non-adhesive areas, therefore controlling their overall shape.

We tested a large number of putative adherent and repulsive coatings, with dose dependent conditions, whose properties were evaluated using the MAAACS algorithm ^42^, a refined cell tracking detection software which includes quantification of intracellular fluorescent signals such as calcium fluxes together with cell shape and speed. We took advantage of this latter parameter as a global readout of the capacity of our panel of candidate molecules to immobilize cells through adhesion and spreading. Aside, we evaluated whether those compounds were prone to activate or to anergize T cells, by recording the calcium response of cells once they reached the surface, followed, when applicable by an additional step of activation with saturating concentration of soluble aCD3.

As reported on Fig. 1, we observed that T cells seeded onto aCD3-coated surface were adhering (i.e. instant speed drops rapidly after contact) and activating strongly (rise of the fluorescence amplitude of the PBX calcium indicator), while on aCD45-coated glass surface (H193.16.3 clone), T cells were adherent as well, spread vigorously without inducing a rise of calcium influx (consistent with our previous observations ^42^) unless subsequently stimulated with soluble aCD3. To the contrary, T cells did not adhere on serum-, PLL-peg-, F127-, F108-, (biotin)-BSA-coated surfaces. On PLL, we highlighted adhesion without spreading which appears to be insufficient to prevent T cells from detaching upon soluble aCD3 stimulation. Biotinylated BSA+streptavidin coating induced for some T cells a slight activation compared to BSA or biotinylated BSA alone on continuous substrates but this coating was essentially non adherent, hence classifying it as repulsive and non activatory. On all other tested conditions (namely fibronectin, mICAM1, WGA, concanavalin A, protein A [without aCD3 or aCD45], aCD43, aMHCI), T cells were either loosely adherent and/or activated by the substrate.

**Figure 1.**
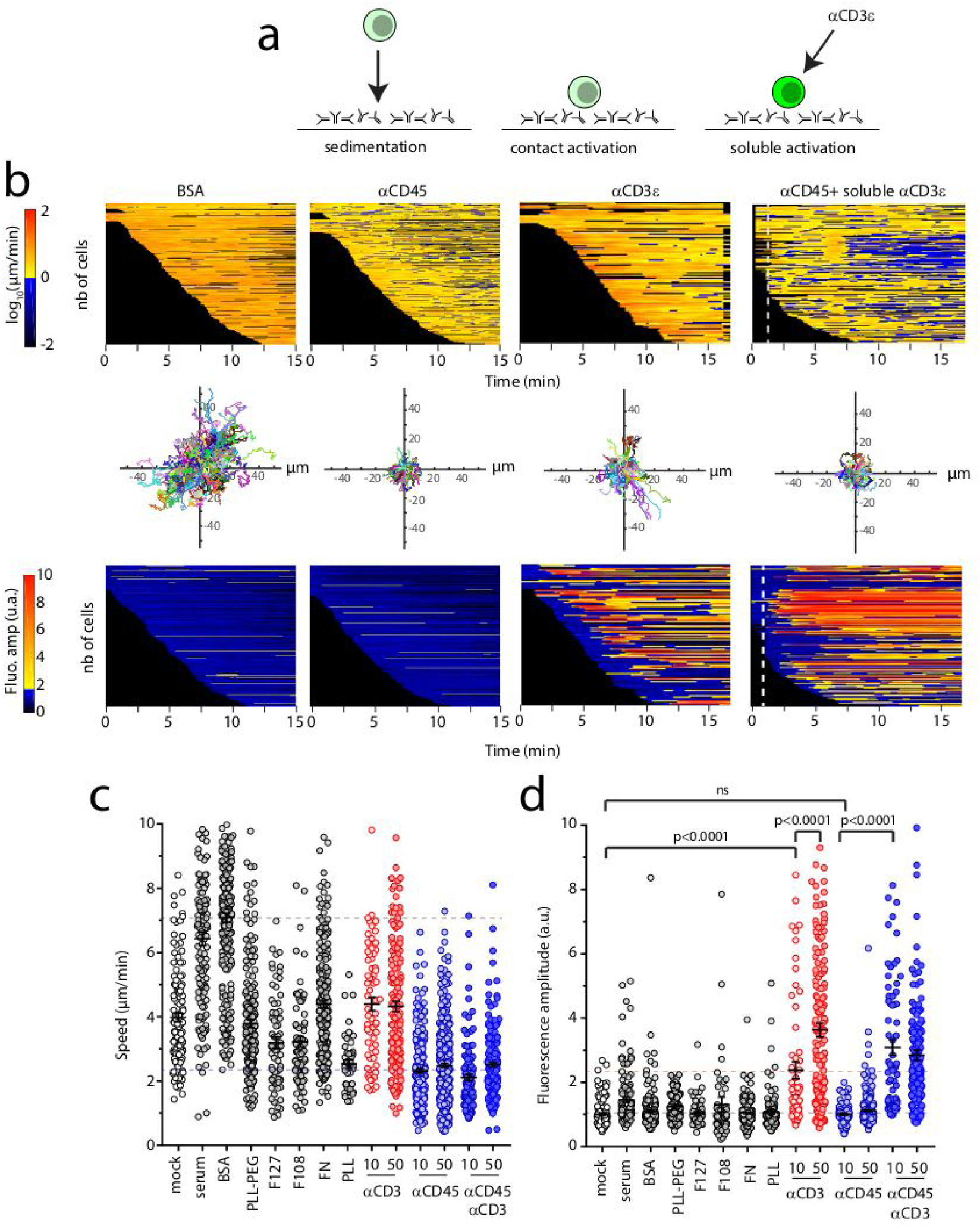
**a**. Schematics of a MAAACS ^42^ experiment, where cells first sediment on a substrate which may be passivated, adherent or activating and are, in some cases, further stimulated with soluble aCD3. **b**. Heatmaps of cell speed, re-aligned cellular trajectories and heatmaps of normalized calcium fluxes vs. time extracted from representative movies on (i) a non adhesive substrate (BSA 1%), (ii) an adhesive but non activating substrate (aCD45), (iii) an adhesive and activating substrate (aCD3), and (iv) on cells after landing on (ii), but subjected to reactivation via soluble aCD3 (the instant of injection is presented as a dotted line). One line corresponds to one detected cell. **c**. Measured speed of cells, from MAAACS (see text), which is used to define the adhesiveness of the tested substrates : when average speed is < 3µm/min, the substrate is defined as adhesive. **d**. Fluorescence amplitude of cells, corresponding to substrates tested in **c**. Data is presented as mean +/- SEM. Significance of difference is assessed with Mann-Whitney test.

Due to its abundance and its large extracellular domain, CD45 has been used by many authors over years as a tool molecule, which can be used to efficiently immobilize T cells without affecting their biology ^61^, although some reports indicate that CD45 segregation may nevertheless impact T cell activation ^62^, depending on the isotype of the antibody used ^63^. Recent reports show that CD45 segregation is not required to stimulate Jurkat T cells by immobilized recombinant monovalent antibodies raised against CD3 ^64^. We tested other antibodies, such as aCD43 or aMHCI following ref. ^25^, but cells were less adherent on such substrates, while eventually showing calcium fluctuations above threshold.

This therefore qualified the H193.16.3 aCD45 antibody as a candidate which fulfills the majority of our initial requirements, even using high concentrations of coated antibody (up to 50µg/ml overnight incubation).

### Adhesive/repulsive micropatterns for T cells shaping

We chose to pattern protein circles since (i) this shape offers the largest adhesive area possible, (ii) it allows for a simple geometry to test from AFM indentation experiments, eg. for placing the indenter on reproducible part of the cell, with a known nucleus position (see text), (iii) it is known that T cells clones tend to become more spherical upon activation^65^ and (iv) all of this will contribute in our effort to minimise data dispersion.

To design our micropatterning strategy, we quantified the spreading area of 3A9m cells on continuously decorated aCD45 substrates vs. time over relevant periods of time (up to 1hr) by RICM (Fig. 2a,b). From the maximal area the 3A9m cells spread, we then calculated an equivalent maximal radius of our circular patterns (2R∼20 µm), hence constraining our designs to smaller sizes. In our hands, patterns smaller than 10µm in diameter were not able to properly immobilize the T cells which would be a major impairment for AFM experiments.

**Figure 2.**
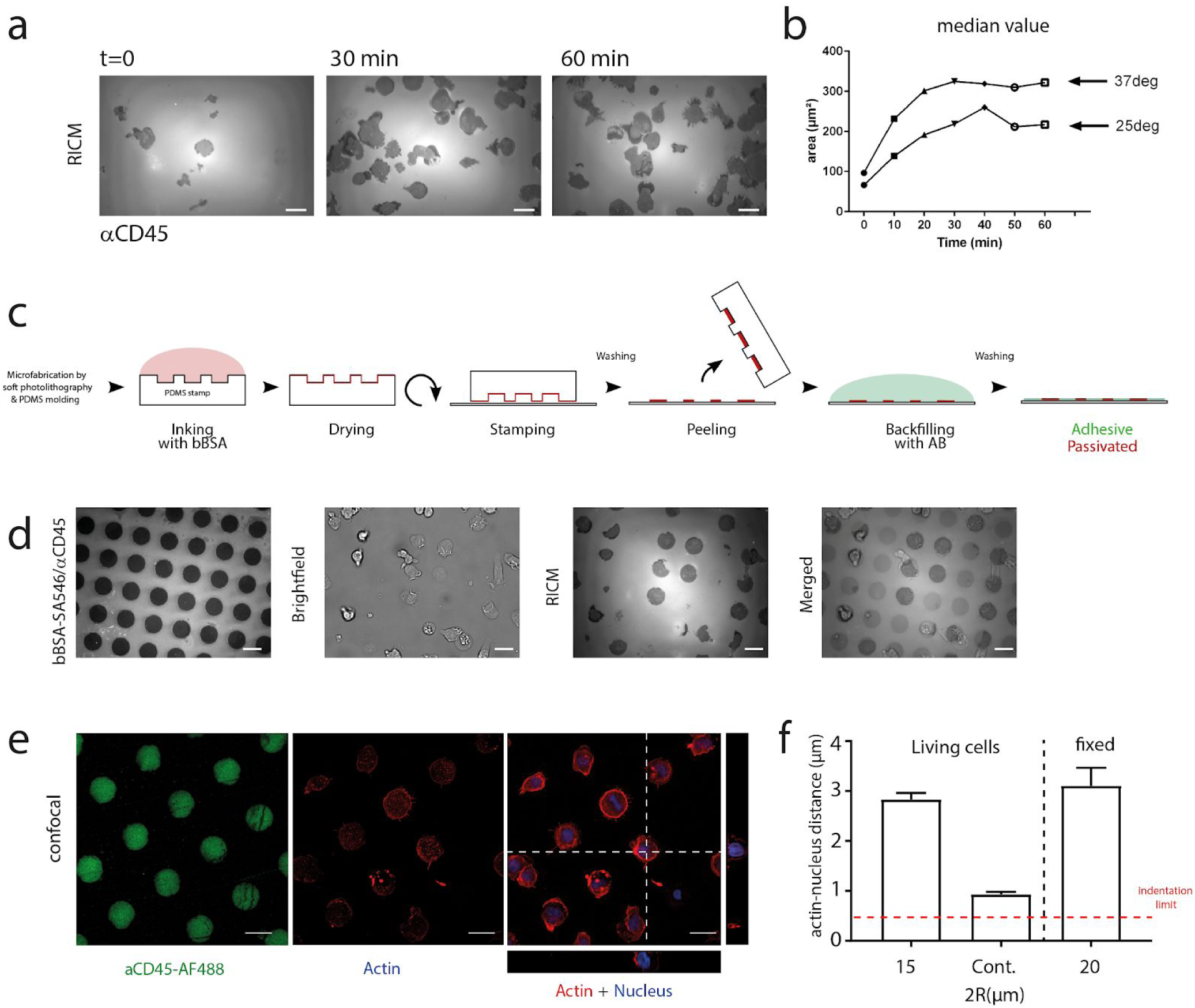
**a**. RICM images of adhesion and spreading of 3A9m cells on aCD45 “continuous” (denominated as “cont.”) substrate at given time points. **b**. Median adhesion area on aCD45 of cells vs. time at 25°C and 37°C. The adhesion after 60 min at 37°C corresponds to an equivalent disk of diameter of ∼19 µm. **c**. Schematics of the inverse microstamping method, where the repulsive molecule (BSA or bBSA) is stamped first, before adsorbing the adhesive one (aCD45) in the free zones. **d**. Image of obtained substrates : (i) fluorescence image of the repulsive zones, completed with non labeled aCD45 in the patterns (dark), (ii) brightfield images of 3A9m cells after 60 min incubation, (iii) corresponding RICM image of the adhesion zones, (iv) overlay of the three previous pictures. **e**. Confocal imaging of 3A9m cells showing patterns of aCD45 (green) cortical actin (red) and nucleus (blue), with projections of the z-stack. **f**. Estimation of the distance between the cortical actin and the nucleus (mean+/SEM), to determine the maximal indentation for the Hertz-like fit (dotted line, see text). Bars are 20µm.

We first successfully directly stamped disks of Alexa Fluor 647 labelled aCD45 and backfilled with either BSA, PLL-PEG or F127. On these substrates, denominated “direct micro-contact printing”, the time for the T cells to spread was rather large, usually more than 1 hour and reproducibility of the spreading was not achieved, the substrates showing often incomplete patterns or even imperfect cellular adhesion as seen in RICM. We concluded that either we were losing a large amount of antibodies while patterning, resulting in the patterns being polluted by the repulsion molecules we backfilled with, or that our antibodies were sensitive to the drying step needed before stamping, hence being partially damaged by the process. Quantifying the fluorescence of aCD45 antibodies stamped to a glass substrate, after incubation on a stamp, to the one of adsorbed antibodies, we estimated that the loss could be ∼ 40-50 % as compared to a simple 45 min adsorption procedure (Suppl. Fig. 1)^66^.

As a consequence, we decided to stamp repulsive molecules first with inverted patterns corresponding to the previous ones, then backfill them by adsorbing aCD45 antibodies. Notably, we did not succeed in transferring efficiently PLL-PEG or pluronic F127, even by modifying either the stamp or the glass substrate using plasma activation or UV exposure. Aside, we obtained good results when the repulsive proteins were BSA or biotinylated BSA onto unmodified or plasma activated glass (Fig. 2b).

We then optimized the protocols to obtain the higher adhesive / repulsive contrast when cells were seeded onto bi-functional substrates. We patterned biotinylated BSA, adsorbed aCD45 and finally functionalized the biotinylated BSA with fluorescent streptavidin. This solution afforded us to use unlabeled aCD45 in conjunction with labelled streptavidin in order to reveal the proper (inverse) patterning of our substrates while avoiding any fluorescence crosstalk for recording calcium probe signals.

To be compatible with our planned mechanical tests using AFM, we wanted to (a) minimize the time needed for T cells to adhere on the larger stamps of aCD45, (b) play on spreading temperature to minimize calcium dye compartmentalization and (c) wash gently the non-adherent cells away while avoiding displacing / perturbing the patterned ones. We determined that an incubation of the cells for 1h at 25°C, when previously pre-loaded with the calcium reporter molecule, allowed a good compromise between criteria (a) and (b). Thus, the protocol presented in the Material and Methods section is the one we observed to be optimal and tractable for our experimental needs.

Cells were spreading circularly as expected on the pattern, exhibited similar areas on the largest patterns and on continuous substrates but different cell morphologies : they were more elongated (4π × *Area*/*Perimeter*^2^ ∼0.6+/-0.2) and flatter (see below) on continuous adherent substrates. They appeared to have a dynamical membrane, but were not motile even on continuous substrates (as recorded by our MAAACS experiments), and could be kept alive up to 3 hrs for non-labelled cells before starting to present vacuoles (not shown), a very interesting point for AFM measurements as described hereafter.

### T cell mechanics of patterned T cells

Cell mechanics was characterized by AFM indentation using bead decorated cantilevers (Fig. 3a,b). We repeatedly indented adhered cells with a moderate maximum contact force, with contact times of 0 sec or up to several seconds. In the first case, we extracted the Young modulus from the contact part of the force curve ^43,58,67^, and in the second case we also extracted the exponent of a power law fit of the force relaxation during the first seconds of contact ^60^.

**Figure 3.**
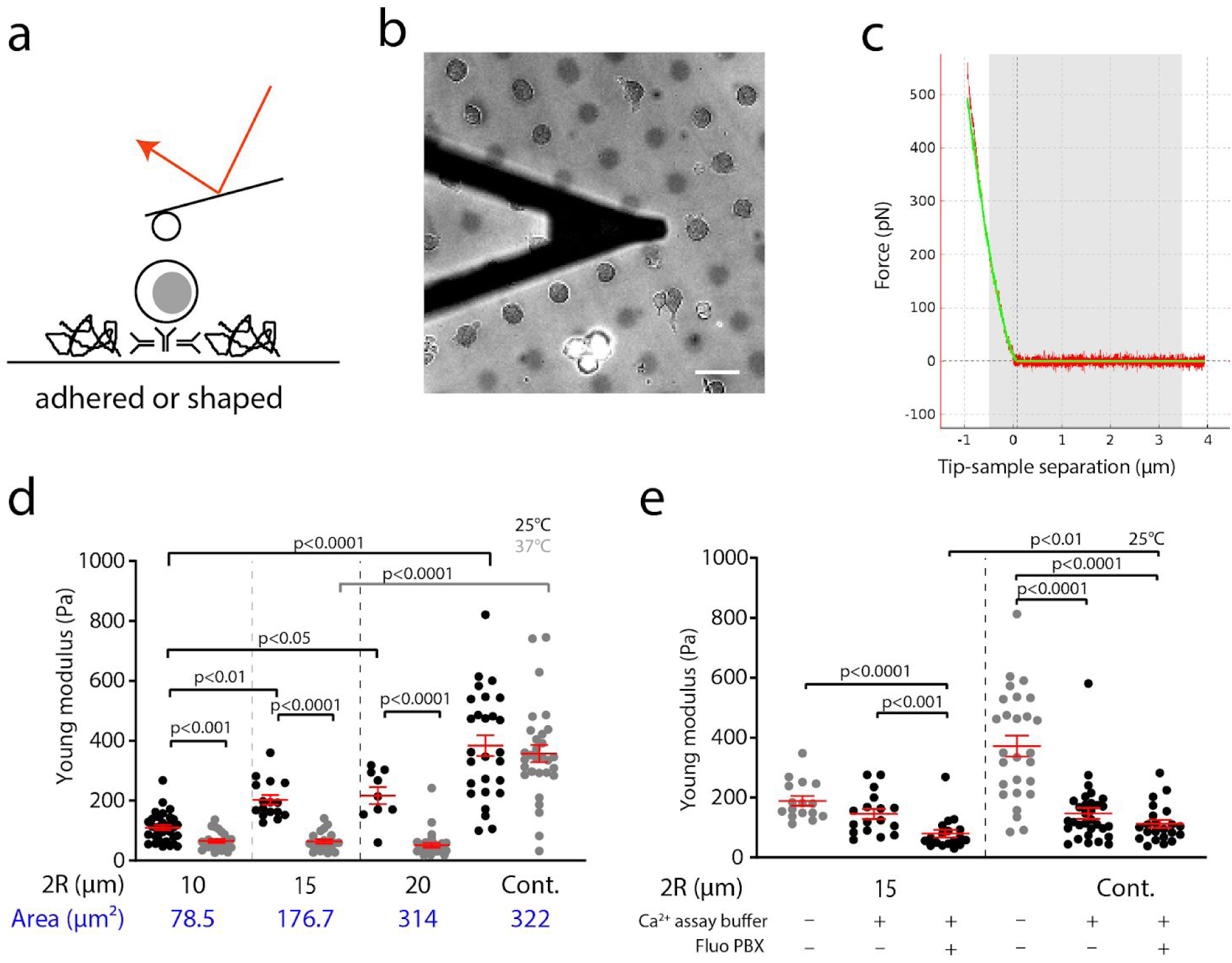
**a**. Schematics of AFM indentation experiments on patterned 3A9m cells. **b**. Micrograph (bar=20µm) of a bead decorated lever (black triangular shadow) close to patterned 3A9m cells. The image is an overlay of the fluorescence of the non adhesive zone and of the corresponding brightfield image. **c**. Representative indentation force curve (only the pushing part), with a Hertz fit (green curve) for a spherical indenter over the first 0.5µm of indentation (grey zone). **d**. Young modulus measurements of patterned and non-patterned cells performed at 25°C and 37°C. **e**. Effect of the incubation of the PBX dye and loading buffer on the measured Young modulus (at 25°C) of the patterned and non-patterned cells. In grey, data from **d** for comparison. **d, e** : One point is the average modulus obtained over up to 10 repeated indentations of the same cell. Red symbols are for mean ± SEM. Significance of difference is assessed with a Mann-Whitney test. Continuous substrate is denominated as “cont.”.

In order to fix the maximal indentation depth for the Hertz fit, we imaged using confocal microscopy the relative positions of the membrane and of the nucleus on adhered cells (Fig.2e,f). We obtained typical values larger than 1µm and consequently chose to fit the experimental indentation curves over the first 0.5µm of indentation to exclude as much as possible any bias due to the nucleus.

Our measurements confirmed that temperature is a crucial parameter for T cell mechanics (Fig. 3c) as for other leukocytes ^68^. When experiments were performed at 25°C, the measured Young modulus gradually increased following the spreading area. In contrast, when the experiments were performed at 37°C, the only marked difference was observed for patterned vs. continuous substrates, the three pattern sizes inducing no visible change or trend in Young modulus. The values we obtained for weakly constrained cells at 25°C or at 37°C for all stamp sizes are coherent with the published values for T cells that are only weakly aspirated in micropipettes, without adhesion, suggesting that our method does not affect importantly T cell mechanics as long as activation is controlled ^66^.

For the patterns, our results point out toward a possible mechanical homeostasis, being efficient at 37°C, for the time during which the sample are prepared ^69^. Interestingly, the difference between patterned and non-patterned cells could be linked to the morphology after spreading, since non patterned cells are flatter than patterned ones : the membrane / nucleus distance was observed to be significantly smaller. (Fig. 2e,f) This may point toward a residual influence of the nucleus on the measured mechanics eg. by differential densities or organisation of (sub-membrane / above the nucleus) actin ^70^, but our confocal imaging did not allow us to resolve such organisation.

The force relaxation power law exponent was weakly sensitive to the modality of immobilization of the cells and the estimated values were, interestingly, on the same order of magnitude of the ones reported for other cell types using similar methodologies ^60^. The median values of the relaxation exponent *n* we estimated were resp. -0.14, -0.17, -0.15, -0.09 for stamps of 10, 15, 20 µm in diameter and continuous substrates, at 37°C, suggesting that the patterned T cell show a similar viscous behavior while the cells spread on continuous substrate are more viscous.

Moreover, the loading of T cells with the calcium probe did not influence neither T cells shape nor adhesion. Surprisingly, their Young modulus was slightly lower than for non-loaded cells of the same shape and spreading. We tracked this perturbation to the commercial loading solution provided with the reporter dye which may contain some surfactants to help the dye permeate and could not rule out that the slight calcium sequestration by the probe may introduce a perturbation of the cytoskeleton architecture ^71^.

### Shaping T cells through aCD45 patterns does not affect their activation by soluble aCD3

Using MAAACS, we observed that soluble aCD3 stimulation of patterned cells (ie. after 1hr spreading at 25°C before returning the cells to 37°C) induced robust calcium signals comparable to the classical MAAACS situation, where T cells were spread on continuous aCD45 substrates (for the same duration) (Fig. 4).

**Figure 4.**
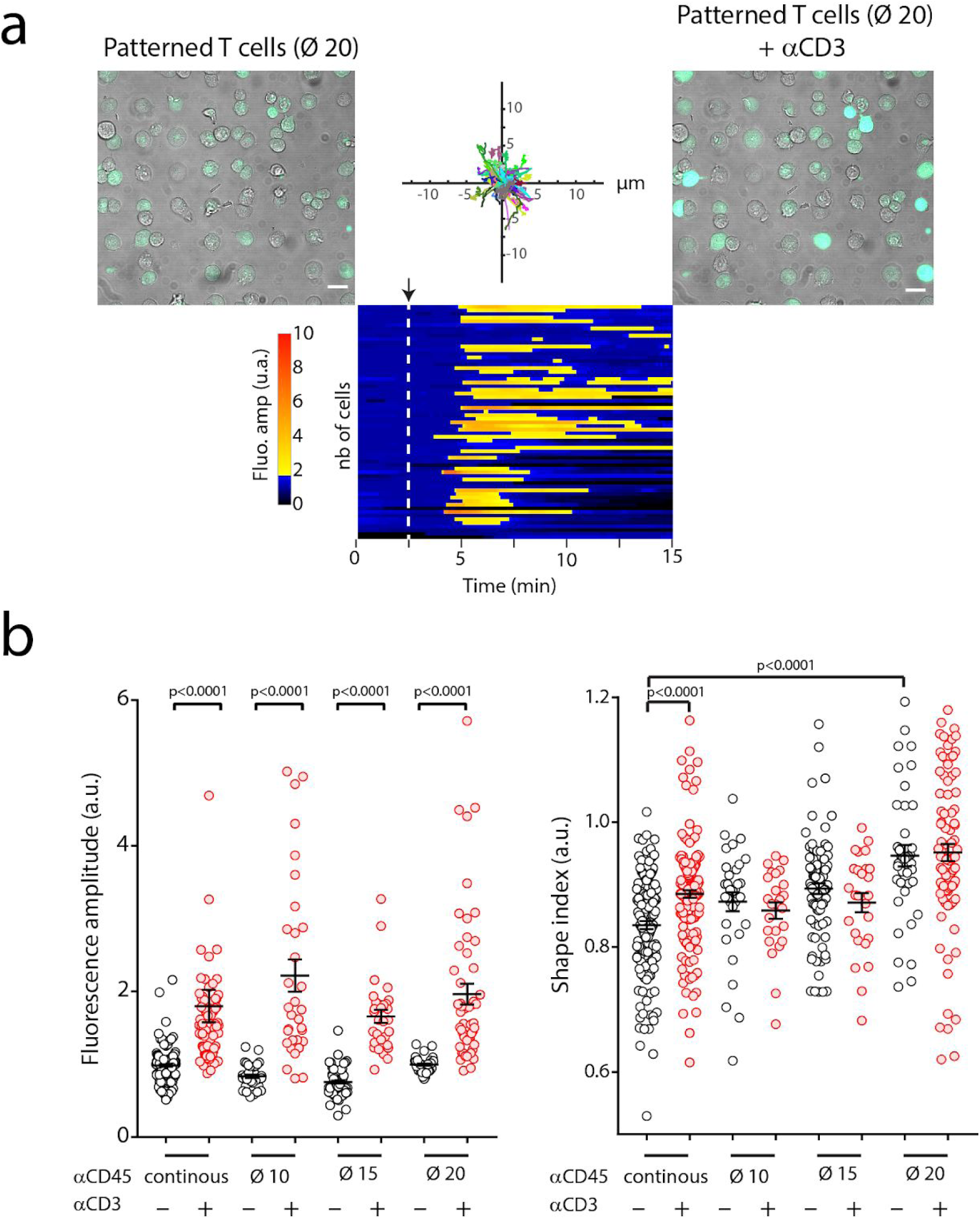
**a**. MAAACS experiments with 3A9m cells sitting 20µm aCD45 patterns. Representative images before and after addition of soluble aCD3. Re-aligned trajectories, showing that cells are essentially immobile. Heatmap of fluorescence amplitude vs. time of only patterned cells, with the instant of aCD3 addition shown with a dashed line. One line is one detected cell. **b**. Fluorescence amplitude before and after injection of aCD3 for cells patterned or not. Shape index variation with the introduction of aCD3, showing that non shape restricted cells are rounding upon activation with soluble aCD3. One point in one cell. Black symbols are for mean +/- SEM. Significance of difference is assessed with a Mann-Whitney test.

Since the mechanical properties of the cells were similar at 37°C among the patterns but not with the continuous situation, and that the largest patterns lead to a same spreading area than the continuous substrate, we propose that, in our conditions, calcium fluxes elicited by soluble stimulation are neither affected by cell’s geometry, cell’s elasticity nor cell anisotropy of spreading.

Interestingly, the amount of CD45 sequestered at the basal side of the cells does not appear to affect neither the intensity nor the dynamics (i.e. the time delay before calcium rise or speed of this rise) : from 10 to 20 µm in diameter for the patterns, the apparent adhesive area is increased by a factor 4, leading to a possible 4-times larger consumption of CD45 molecules.

### Ectopic expression of an non-immunologic adhesion molecule

To further verify that the impact of CD45 was minimal in our system, we ectopically expressed in 3A9m cells an adhesion molecule that is not expressed in T cells. We chose the MEGF10 protein (for Multiple Epidermal Growth Factor-like domain Protein 10, Fig. 5a, ^49^). This protein is the orthologue of the Ced-1 receptor involved in the recognition of apoptotic cells in the nematode C. Elegans ^72^. Various studies in Drosophila (DRAPER gene) or in higher eukaryotes have shown that this family of membrane proteins has retained this function ^73^. However, it can be noted that the expression of MEGF10 in mammals is restricted to the cerebral (expressed in astrocytes ^74^, in retinal cells ^75^) and muscular spheres where MEGF10 would influence the proliferation of muscle cells ^76^.

**Figure 5.**
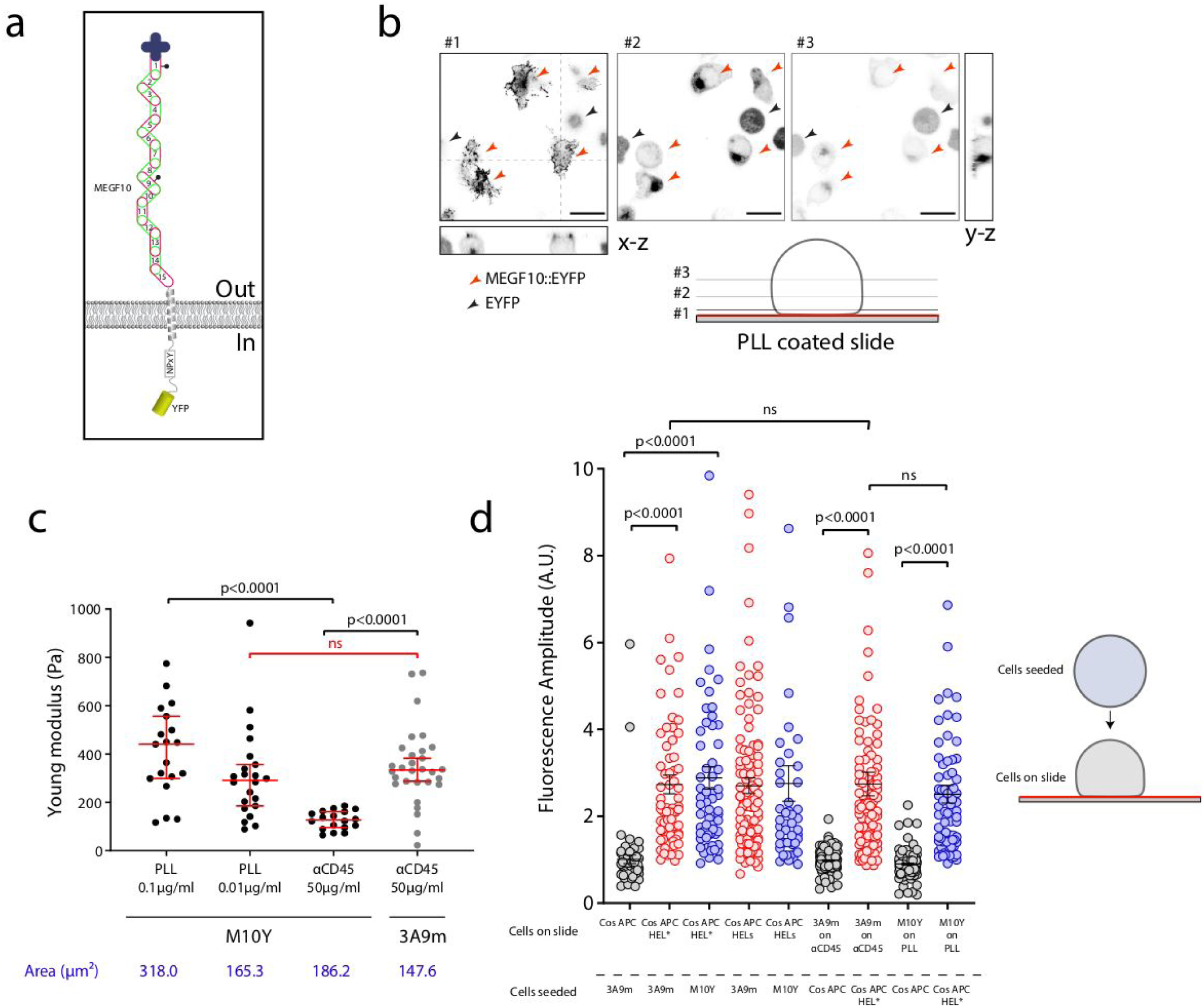
**a**. Schematics of the structure of MEGF10 construct as a membrane receptor, labelled with YFP, expressed in 3A9m M10Y. **b**. Confocal imaging of the polarised MEGF10 localisation when the cells are adhering on PLL substrate, showing that MEGF10 is mainly segregated at the contact zone(#1-3 confocal sections at various altitudes z); red arrows: M10Y expressing 3A9m T cells; black arrow : cytosolic YFP expressing 3A9m T cells (scale bar=20µm). **c**. Variation of the adhesion area and mechanical properties of the transfected cells as compared to aCD45 continuous substrates case. For an intermediate concentration of PLL during the incubation, M10Y cells achieve to spread like WT cells on aCD45 and possess the same mechanical properties. **d**. Activation of M10Y vs. 3A9m cells when seeded on COS APC cells as in classical MAAACS experiments or when adhered respectively on PLL or aCD45 and contacted by sedimenting COS APCs without antigen or I-Ak-HEL expressing COS-7. Results are expressed as mean±SEM and statistical analysis was performed using a Mann-Whitney test.

We electroporated the original MEGF10::YFP construction ^49^ and selected a population of 3A9m cells expressing a comparable level of TCR as original 3A9m and a high level of YFP fluorescence (3A9m M10Y). We obtained a polyclonal population of mixed cells that express either MEGF10::YFP (M10Y+) or only cytosolic YFP (due to a truncated integration of the construction and a shift of an initiating codon at the ATG level of the YFP) (Fig. 5b). However, this M10Y+ population demonstrated a very strong adhesion to poly-L-Lysine (while very weakly adherent on glass alone), and an enrichment of the MEGF10::YFP protein at the basal membrane in contact with the support, in contrast to cytosolic YFP expressing 3A9m T cells that loosely adhere without spreading on this substrate (consistent with observations for 3A9m T cells in a previous section). The M10Y+ T cells spread very quickly (between 5 and 10 minutes) in the form of a dotted veil with local enrichments as spikes, and we noticed the additional presence of small filopod-type structures (Fig. 5b), consistent with the literature ^77^. In comparison, cells expressing only cytosolic YFP did not show this enrichment or membrane extensions (Fig. 5b).

Varying the concentration of poly-L-Lysine, we observed that a high concentration of PLL (0.1%) leads to a greater spread (almost a factor 2 on the area determined by RICM) compared to a concentration of 0.01% (Fig. 5c). In this case, the cell is more rigid, as shown by a higher Young’s modulus. On the other hand, the adhesion of the cells on a coated surface with a conventional concentration of poly-L-Lysine (0.01%) induces adhesion area (as measured by RICM) as well as Young modulus values of the same order as those measured for 3A9m spread on aCD45 (Fig. 5c).

Interestingly when spread on continuous aCD45 substrates, the adhesion area of M10Y+ T cells appears to be close to the one of 3A9m cells, but these cells are softer than all other spreading conditions. One can only hypothesize that, when MEGF10 is not used for adhesion and seggregates to the contact zone, it may perturb the cytoskeleton organization and / or mechanical measurements due to its large extracellular domain.

### Immobilizing T cells strategies do not perturb their activation by APC

We investigated whether T cells made adherent onto functionalized surfaces would be still activated by antigen presenting cells. To that purpose, we used surrogate APC, namely COS 7 cells expressing the α chain of MHC II and the *β* chain of MHC II I-A^K^ alone (COS APC) or covalently fused to the a peptide derived from the Hen Egg Lysozyme HEL (COS APC HEL*) ^42,48,50,78^. Alternatively COS APC were loaded with soluble antigenic peptide (aa 48-63) derived from HEL (COS APC HEL).

We performed two symmetrical experiments. First, the 3A9m T cells M10Y+ were adhered to the poly-L-Lysine coated surface, then a suspension of COS APCs was added and let fall by gravity on the spread T cells on continuous substrates. For comparison, 3A9m cells were seeded on surfaces covered with aCD45 antibodies. Conditions were matched to have cells spread similarly, hence having the same mechanical properties (see above). Second, conversely, COS APC cells and variants were adhered at the bottom of the observation chambers, and 3A9m or M10Y cells being seeded over them in a similar way as previously ^42,50^. In both cases, the T cells have been pre-loaded with PBX calcium reporter, and experiments performed at 37°C.

The percentage of activated cells in the configuration where the T cells are adherent is generally lower than in the reverse situation (immobile APC and sedimenting / mobile T cells). This difference can be explained by the T/APC cell ratio since a good number of T cells present in the field of observation were not in contact by seeded Cos APC cells. Following the calcium fluxes, we observed that the fluorescence amplitude of M10Y+ T cells (on PLL; FA =1.96±0.20) is highly similar to than the 3A9m adhered to on aCD45 (FA=1.73±0.11; p=0.498, Mann-Whitney test). Globally, whatever the configuration chosen or the experimental conditions, we could not find any significant difference between the activation signature of 3A9m cells and the M10Y+ T cells by model APCs (Fig 5d).

These observations suggest that at equivalent expression level of TCR, immobilizing CD45 on wild type 3A9m T cells does not impact significantly the antigen-dependant calcium release compared to M10Y+ T cells

In addition, we tried to obtain patterned surfaces with PPL as an adhesive molecule, by a direct printing strategy using modified stamps. It appears that in our hands, and with the glass bottom Petri dishes we used to be fully compatible with AFM measurements at 37°C, the transfer of PLL onto the surface is not homogeneous (as seen using with a 9:1 mix of PLL:Alexa Fluor 546-labelled PLL), some part remaining on the PDMS stamp. We have plasma treated the surfaces of glass petri dishes which slightly improved the quality and reproducibility of the transfer. We also tried to first incubate PDMS stamps with 10% Sodium Dodecyl Sulfate (SDS) before inking, since it was shown that such pretreatment may increase the proportion of transferred PLL onto an activated glass surface ^79^.

Moreover, it unfortunately proved to be difficult to obtain functional patterns, suitable for the immobilization of M10Y cells since they also tend to adhere relatively strongly to bare glass and also, surprisingly, BSA so we tested in particular PLL/PLL-Peg combinations, PLL-PEG being used as repulsive coating molecule to backfill the free space between the stamps (Suppl. Fig. 2). Incomplete patterns were observed without SDS pre-treatment of the PDMS stamp, which lead either to non-specific adhesion, with cells adhering everywhere, or to specific but rarely complete adhesion since repulsive molecules were polluting the stamps. In the case of stamps pre-coated with SDS, the cells did not spread over the entire surface of the pattern, indicating a potential contamination with the repulsive molecules.

These limitations, in parallel with our results on the mechanics and activability of the cells, has confirmed our idea that the H193.16.3 CD45 antibody is, in our hands, one of the simpler and better ways to shape murine T cells onto patterns.

## Conclusions and perspectives

Here we present a strategy aiming at controlling T cell shape and mechanical properties, while maintaining a good activability. We used model murine hybridoma T cells which recapitulates the essential steps of T cell activation up to IL-2 production. Using a refined image processing method, named MAAACS, we identified several molecules which could be used to immobilize cells on a glass substrate, and several preventing this adhesion, of which the couple (CD45, BSA/biotin BSA) was chosen. We determined by interference microscopy recording the maximal spreading of the cells on continuous substrate.

We tested direct micro-contact printing of the CD45 and observed that reproducibility of the adhesive behavior was non ensured, and attributed this to a potential denaturation of the antibody during the necessary drying step of our procedure. We then designed an “inverse” printing protocol allowing us to first shape the repulsion, then adsorb the adhesive molecule, and reveal the repulsion using labelled streptavidin, avoiding any labelled molecule to interfere with calcium fluxes detection on the shaped cells. Patterning the repulsion and building the adhesive zones by adsorption also helped us ensure that the amount of CD45 is the same as on continuous substrates, the extension of the adhesive zones allowing us to control how many molecules we propose to the cell to adhere on to test the influence of CD45 on our activation assay.

We then characterized cell mechanics by AFM indentation with moderate forces and noted that spreading on circular patterns of CD45 does not influence cell elasticity. In contrast, asymmetrical cells obtained on continuous substrates, with a comparable adhesion area as for the largest patterns, are more rigid, more likely due to a different positioning of the nucleus. We observed that temperature, but also preliminary incubation with the loading buffer of the calcium dye, has an effect on cell mechanics, with Young modulus being the lower when the solution was used and the cells measured at physiological temperature. The dye itself, even if chelating some of the cytosolic calcium, had only a mild effect on the elasticity.

When tested for activation behavior with aCD3 in solution, patterned cells and non-patterned cells exhibited the same capabilities, indicating that in our assays CD45 consumption by adhesion was not having a strong impact. The key point was to allow the T cell spreading on patterns while maintaining a level of calcium dye relocalization from the cytosol as low as possible by a careful play on the sample temperature.

To rule out that partial CD45 immobilisation would affect T cell activation, we generated a new cell line, expressing a non immunological adhesion molecule that allowed strong adhesion to PLL substrates, compared to wild type 3A9m, even if PLL is not unequivocally considered as the ideal experimental substrate for lymphocyte adhesion ^80^. MEGF10 was shown to localise at the contact between cells and substrates. By tuning the amount of PLL used to decorate the substrate, we found a condition where the spreading area together with the Young modulus of this new cell line were similar to the ones of the original cell line when spread on continuous aCD45.

We verified that, when challenged by model APCs, the two cell lines immobilized on aCD45 or PLL (resp.) were exhibiting the same activation capabilities, underlying the fact that in our experiments, the use of CD45 as an anchor to the surface does not affect activation patterns and intensity as measured using calcium fluxes reporters.

The “platform” of the calcium probe loaded and shaped T cell we designed, with controlled mechanical properties, opens up new possibilities for single molecule approaches such as Single Cell Force Spectroscopy for excluding shape and mechanical effects. It also permits more complex mechanical characterization of the impact of soluble molecules on cell membrane or cytoskeleton, such as by pulling tethers with optical tethers where it has been hypothesized that initial state of the cell influences measurements, and potentially opens up way to study T cell mechanics modulation by controlled surface activation signals (density, amount, shape). It also provides periodic, calibrated T cells for refined imaging such as PALM/STORM or diffusion measurements such as Fluorescence Correlation spectroscopy. By varying T cell shape using asymmetrical patterns, more migrating like shapes of cells could be obtained, allowing to study the impact of cell shape on cell function in a reliable manner.

## Acknowledgements

*Fundings* : PEPS “Innovation frugale” (2017, to PHP), ANR JCJC « DissecTion » (2009-2012, to LL & PHP), Labex INFORM (ANR-11-LABX-0054 and A*MIDEX project (ANR-11-IDEX-0001-02), funded by the « Investissements d’Avenir » French Government program managed by the French National Research Agency (ANR), to Inserm U1067 Lab & CIML, and as PhD grant to AS, 2014-2018). This work was supported by the GDR ImaBio through master’s internships funding (LGD, AW). ImaBio is a national thematic research network structured by the CNRS and centered around biophotonics. http://imabio-cnrs.fr. We acknowledge the PICSL imaging facility of the CIML (ImagImm), member of the national infrastructure France-BioImaging supported by the French National Research Agency (ANR-10-INBS-04) and the CIML cytometry facility)

*Providing material or technical help*: F. Bedu (PLANETE clean room facility, CINaM), L. Borge (PCC cell culture facility), A. Formisano & H.-T. He (CIML), S. Mailfert / M. Fallet / R. Fabre (CIML Imagimm Imaging facility), N. Bertaux (Ecole Centrale Marseille), T. Sbarrato, L. Aoun (LAI), AF. Rigato, F. Rico (Inserm U1067, Marseille).

*Companies*: F. Eghiaian, T. Müller, T. Plake, J. Beaumale & JPK Instruments/Bruker (Berlin, Germany) for continuous support and generous help. Zeiss France for support.

## Contributions

AS performed experiments, processed the data and wrote the article. MBP performed FACS experiments and related analysis. LL participated in data analysis. OT provided microstamps and helped with troubleshooting stamping strategies. LGD and AW participated in optimizing COS cell immobilization and T cell stamping strategies. YH and PHP designed, performed experiments, processed the data and wrote the article.

## References

1. Ehrich, E. W. et al. T cell receptor interaction with peptide/major histocompatibility complex (MHC) and superantigen/MHC ligands is dominated by antigen. J Exp Med 178, 713–22 (1993).

2. Purbhoo, M. a, Irvine, D. J., Huppa, J. B. & Davis, M. M. T cell killing does not require the formation of a stable mature immunological synapse. Nat. Immunol. 5, 524–30 (2004).

3. Bromley, S. K. et al. The immunological synapse. Annu Rev Immunol 19, 375–396 (2001).

4. Alam, S. M. et al. T-cell-receptor affinity and thymocyte positive selection. Nature 381, 616–20 (1996).

5. Goldrath, A. W. & Bevan, M. J. Selecting and maintaining a diverse T-cell repertoire. Nature 402, 255–62 (1999).

6. Goldrath, A. W. & Bevan, M. J. Low-affinity ligands for the TCR drive proliferation of mature CD8+ T cells in lymphopenic hosts. Immunity 11, 183–90 (1999).

7. Utzny, C., Faroudi, M. & Valitutti, S. Frequency Encoding of T-Cell Receptor Engagement Dynamics in Calcium Time Series. Biophys. J. 88, 1–14 (2005).

8. Schamel, W. W. et al. Coexistence of multivalent and monovalent TCRs explains high sensitivity and wide range of response. J Exp Med 202, 493–503 (2005).

9. Weiss, A. & Littman, D. R. Signal transduction by lymphocyte antigen receptors. Cell 76, 263–74 (1994).

10. Germain, R. N. The T cell receptor for antigen: signaling and ligand discrimination. J Biol Chem 276, 35223–6. (2001).

11. van der Merwe, P. A. & Dushek, O. Mechanisms for T cell receptor triggering. Nat. Rev. Immunol. 11, 47–55 (2011).

12. Gil, D., Schamel, W. W., Montoya, M., Sanchez-Madrid, F. & Alarcon, B. Recruitment of Nck by CD3 epsilon reveals a ligand-induced conformational change essential for T cell receptor signaling and synapse formation. Cell 109, 901–12 (2002).

13. Krogsgaard, M. et al. Agonist/endogenous peptide-MHC heterodimers drive T cell activation and sensitivity. Nature 434, 238–43 (2005).

14. Rosette, C. et al. The impact of duration versus extent of TCR occupancy on T cell activation: a revision of the kinetic proofreading model. Immunity 15, 59–70 (2001).

15. Huang, J. et al. The kinetics of two-dimensional TCR and pMHC interactions determine T-cell responsiveness. Nature 464, 932–6 (2010).

16. Axmann, M., Huppa, J. B., Davis, M. M. & Schütz, G. J. Determination of interaction kinetics between the T cell receptor and peptide-loaded MHC class II via single-molecule diffusion measurements. Biophys. J. 103, L17–9 (2012).

17. Robert, P. et al. Kinetics and Mechanics of Two-Dimensional Interactions between T Cell Receptors and Different Activating Ligands. Biophys. J. 102, 248–257 (2012).

18. Limozin, L. & Puech, P.-H. Membrane Organization and Physical Regulation of Lymphocyte Antigen Receptors: A Biophysicist’s Perspective. J. Membr. Biol. 252, 397–412 (2019).

19. He, H.-T. & Bongrand, P. Membrane dynamics shape TCR-generated signaling. Front. Immunol. 3, 90 (2012).

20. Malissen, B. & Bongrand, P. Early T Cell Activation: Integrating Biochemical, Structural, and Biophysical Cues. Annu. Rev. Immunol. 33, 539–561 (2015).

21. Chen, W. & Zhu, C. Mechanical regulation of T-cell functions. Immunol. Rev. 256, 160–176 (2013).

22. Puech, P.-H. et al. Force Measurements of TCR/pMHC Recognition at T Cell Surface. PLoS ONE 6, e22344 (2011).

23. Liu, B., Chen, W., Evavold, B. D. & Zhu, C. Accumulation of dynamic catch bonds between TCR and agonist peptide-MHC triggers T cell signaling. Cell 157, 357–368 (2014).

24. Limozin, L. et al. TCR-pMHC kinetics under force in a cell-free system show no intrinsic catch bond, but a minimal encounter duration before binding. Proc. Natl. Acad. Sci. U. S. A. 116, 16943–16948 (2019).

25. Brodovitch, A., Bongrand, P. & Pierres, A. T lymphocytes sense antigens within seconds and make a decision within one minute. J. Immunol. Baltim. Md 1950 191, 2064–2071 (2013).

26. Brodovitch, A. et al. T lymphocytes need less than 3 min to discriminate between peptide MHCs with similar TCR-binding parameters. Eur. J. Immunol. n/a-n/a (2015) doi: 10.1002/eji.201445214.

27. Kim, S. T. et al. The alphabeta T cell receptor is an anisotropic mechanosensor. J Biol Chem 284, 31028–37 (2009).

28. Kim, S. T. et al. TCR Mechanobiology: Torques and Tunable Structures Linked to Early T Cell Signaling. Front. Immunol. 3, 76 (2012).

29. Li, Y.-C. et al. Cutting Edge: mechanical forces acting on T cells immobilized via the TCR complex can trigger TCR signaling. J. Immunol. Baltim. Md 1950 184, 5959–5963 (2010).

30. Hu, K. H. & Butte, M. J. T cell activation requires force generation. J. Cell Biol. 213, 535–542 (2016).

31. Thauland, T. J., Hu, K. H., Bruce, M. A. & Butte, M. J. Cytoskeletal adaptivity regulates T cell receptor signaling. Sci. Signal. 10, eaah3737 (2017).

32. Judokusumo, E., Tabdanov, E., Kumari, S., Dustin, M. L. & Kam, L. C. Mechanosensing in T lymphocyte activation. Biophys. J. 102, L5–7 (2012).

33. Saitakis, M. et al. Different TCR-induced T lymphocyte responses are potentiated by stiffness with variable sensitivity. eLife 6, (2017).

34. Bufi, N. et al. Human Primary Immune Cells Exhibit Distinct Mechanical Properties that Are Modified by Inflammation. Biophys. J. 108, 2181–2190 (2015).

35. Wahl, A. et al. Biphasic mechanosensitivity of T cell receptor-mediated spreading of lymphocytes. Proc. Natl. Acad. Sci. U. S. A. 116, 5908–5913 (2019).

36. Neves, S. R. et al. Cell Shape and Negative Links in Regulatory Motifs Together Control Spatial Information Flow in Signaling Networks. Cell 133, 666–680 (2008).

37. Negulescu, P. A., Krasieva, T. B., Khan, A., Kerschbaum, H. H. & Cahalan, M. D. Polarity of T cell shape, motility, and sensitivity to antigen. Immunity 4, 421–430 (1996).

38. Anvari, B., Torres, J. H. & McIntyre, B. W. Regulation of pseudopodia localization in lymphocytes through application of mechanical forces by optical tweezers. J. Biomed. Opt. 9, 865–872 (2004).

39. Comrie, W. A., Babich, A. & Burkhardt, J. K. F-actin flow drives affinity maturation and spatial organization of LFA-1 at the immunological synapse. J. Cell Biol. 208, 475–491 (2015).

40. Comrie, W. A., Li, S., Boyle, S. & Burkhardt, J. K. The dendritic cell cytoskeleton promotes T cell adhesion and activation by constraining ICAM-1 mobility. J. Cell Biol. 208, 457–473 (2015).

41. Comrie, W. A. & Burkhardt, J. K. Action and Traction: Cytoskeletal Control of Receptor Triggering at the Immunological Synapse. Front. Immunol. 7, (2016).

42. Salles, A. et al. Barcoding T Cell Calcium Response Diversity with Methods for Automated and Accurate Analysis of Cell Signals (MAAACS). PLoS Comput. Biol. 9, e1003245 (2013).

43. Cazaux, S. et al. Synchronizing atomic force microscopy force mode and fluorescence microscopy in real time for immune cell stimulation and activation studies. Ultramicroscopy 160, 168–181 (2016).

44. Hosseini, B. H. et al. Immune synapse formation determines interaction forces between T cells and antigen-presenting cells measured by atomic force microscopy. Proc. Natl. Acad. Sci. U. S. A. 106, 17852–17857 (2009).

45. Hoffmann, S. et al. Single cell force spectroscopy of T cells recognizing a myelin-derived peptide on antigen presenting cells. Immunol. Lett. 136, 13–20 (2011).

46. Lim, T. S., Mortellaro, A., Lim, C. T., Hämmerling, G. J. & Ricciardi-Castagnoli, P. Mechanical interactions between dendritic cells and T cells correlate with T cell responsiveness. J. Immunol. Baltim. Md 1950 187, 258–265 (2011).

47. Thery, M. & Piel, M. Adhesive Micropatterns for Cells: A Microcontact Printing Protocol. Cold Spring Harb. Protoc. 2009, pdb.prot5255-pdb.prot5255 (2009).

48. Carson, R. T., Vignali, K. M., Woodland, D. L. & Vignali, D. a a. T cell receptor recognition of MHC class II-bound peptide flanking residues enhances immunogenicity and results in altered TCR V region usage. Immunity 7, 387–399 (1997).

49. Hamon, Y. et al. Cooperation between engulfment receptors: the case of ABCA1 and MEGF10. PloS One 1, e120 (2006).

50. Chouaki-Benmansour, N. et al. Phosphoinositides regulate the TCR/CD3 complex membrane dynamics and activation. Sci. Rep. 8, 4966 (2018).

51. Limozin, L. & Sengupta, K. Quantitative reflection interference contrast microscopy (RICM) in soft matter and cell adhesion. Chemphyschem Eur. J. Chem. Phys. Phys. Chem. 10, 2752–68 (2009).

52. Dillard, P., Varma, R., Sengupta, K. & Limozin, L. Ligand-mediated friction determines morphodynamics of spreading T cells. Biophys. J. 107, 2629–2638 (2014).

53. Robert, P., Sengupta, K., Puech, P.-H., Bongrand, P. & Limozin, L. Tuning the formation and rupture of single ligand-receptor bonds by hyaluronan-induced repulsion. Biophys J 95, 3999–4012 (2008).

54. Schindelin, J. et al. Fiji: an open-source platform for biological-image analysis. Nat. Methods 9, 676–682 (2012).

55. Butt, H.-J. & Jaschke, M. Calculation of thermal noise in atomic force microscopy. Nanotechnology 6, 1 (1995).

56. Edelstein, A. D. et al. Advanced methods of microscope control using µManager software. J. Biol. Methods 1, e10 (2014).

57. Hammadi, M., Delcroix, V., Vacher, A.-M., Ducret, T. & Vacher, P. CD95-Mediated Calcium Signaling. Methods Mol. Biol. Clifton NJ 1557, 79–93 (2017).

58. Sadoun, A. & Puech, P.-H. Quantifying CD95/cl-CD95L Implications in Cell Mechanics and Membrane Tension by Atomic Force Microscopy Based Force Measurements. Methods Mol. Biol. Clifton NJ 1557, 139–151 (2017).

59. Sawicka, A. et al. Micropipette force probe to quantify single-cell force generation: application to T-cell activation. Mol. Biol. Cell 28, 3229–3239 (2017).

60. Guillou, L., Babataheri, A., Puech, P.-H., Barakat, A. I. & Husson, J. Dynamic monitoring of cell mechanical properties using profile microindentation. Sci. Rep. 6, (2016).

61. Sherman, E. et al. Article Functional Nanoscale Organization of Signaling Molecules Downstream of the T Cell Antigen Receptor. Immunity 35, 705–720 (2011).

62. Cordoba, S. P. et al. The large ectodomains of CD45 and CD148 regulate their segregation from and inhibition of ligated T-cell receptor. Blood 121, 4295–4302 (2013).

63. Goldman, S. J. et al. Differential activation of phosphotyrosine protein phosphatase activity in a murine T cell hybridoma by monoclonal antibodies to CD45. J. Biol. Chem. 267, 6197–6204 (1992).

64. Al-Aghbar, M. A., Chu, Y.-S., Chen, B.-M. & Roffler, S. R. High-Affinity Ligands Can Trigger T Cell Receptor Signaling Without CD45 Segregation. Front. Immunol. 9, 713 (2018).

65. Donnadieu, E., Bismuth, G. & Trautmann, A. Antigen recognition by helper T cells elicits a sequence of distinct changes of their shape and intracellular calcium. Curr. Biol. CB 4, 584–595 (1994).

66. Guillou, L. et al. T-lymphocyte passive deformation is controlled by unfolding of membrane surface reservoirs. Mol. Biol. Cell 27, 3574–3582 (2016).

67. Franz, C. M. & Puech, P.-H. Atomic Force Microscopy: A Versatile Tool for Studying Cell Morphology, Adhesion and Mechanics. Cell. Mol. Bioeng. 1, 289–300 (2008).

68. Rico, F., Chu, C., Abdulreda, M. H., Qin, Y. & Moy, V. T. Temperature modulation of integrin-mediated cell adhesion. Biophys. J. 99, 1387–1396 (2010).

69. Webster, K. D., Ng, W. P. & Fletcher, D. A. Tensional Homeostasis in Single Fibroblasts. Biophys. J. 107, 146–155 (2014).

70. Versaevel, M., Grevesse, T. & Gabriele, S. Spatial coordination between cell and nuclear shape within micropatterned endothelial cells. Nat. Commun. 3, 671 (2012).

71. Richelme, F., Benoliel, A.-M. & Bongrand, P. Dynamic study of cell mechanical and structural responses to rapid changes of calcium level. Cell Motil. Cytoskeleton 45, 93–105 (2000).

72. Zhou, Z., Hartwieg, E. & Horvitz, H. R. CED-1 Is a Transmembrane Receptor that Mediates Cell Corpse Engulfment in C. elegans. Cell 104, 43–56 (2001).

73. Manaka, J. et al. Draper-mediated and Phosphatidylserine-independent Phagocytosis of Apoptotic Cells by Drosophila Hemocytes/Macrophages. J. Biol. Chem. 279, 48466–48476 (2004).

74. Chung, W.-S. et al. Astrocytes mediate synapse elimination through MEGF10 and MERTK pathways. Nature 504, 394–400 (2013).

75. Kay, J. N., Chu, M. W. & Sanes, J. R. MEGF10 and MEGF11 mediate homotypic interactions required for mosaic spacing of retinal neurons. Nature 483, 465–469 (2012).

76. Logan, C. V. et al. Mutations in MEGF10, a regulator of satellite cell myogenesis, cause early onset myopathy, areflexia, respiratory distress and dysphagia (EMARDD). Nat. Genet. 43, 1189–1192 (2011).

77. Suzuki, E. & Nakayama, M. MEGF10 is a mammalian ortholog of CED-1 that interacts with clathrin assembly protein complex 2 medium chain and induces large vacuole formation. Exp. Cell Res. 313, 3729–3742 (2007).

78. Xia, F. et al. TCR and CD28 Concomitant Stimulation Elicits a Distinctive Calcium Response in Naive T Cells. Front. Immunol. 9, 2864 (2018).

79. Chang, J. C., Brewer, G. J. & Wheeler, B. C. A modified microstamping technique enhances polylysine transfer and neuronal cell patterning. Biomaterials 24, 2863–2870 (2003).

80. Santos, A. M. et al. Capturing resting T cells: The perils of PLL correspondence. Nat. Immunol. 19, 203–205 (2018).

