## Supplementary material for "Controlling T cells shape, mechanics and activation by micropatterning": Suppl. Figs.

**Supplementary figure captions**

**Suppl. Fig 1 :** Comparison of the transfer efficiency of direct vs. inverse microcontact printing techniques on glass bottom Petri dishes with aCD45 coupled to Atto 565, in regard to simple adsorption, as a function of the solution concentration. Duration of incubation between inverse MCP and adsorption were the same (see text). Exponential fits to the data were plotted as a guide for the eyes when more than three data points were available.

**Suppl. Fig 2 :** Stamping 9:1 PLL:Alexa Fluor 546-labelled PLL using PDMS stamps, pre-treated or not with SDS. Backfilling was performed using PLL-PEG. When no SDS was used, the transfer of patterns was poorly resolved. When SDS was used and M10Y cells seeded , the cells failed to completely fill the patterns which were better resolved. Scale bar=20μm.

Figure S1

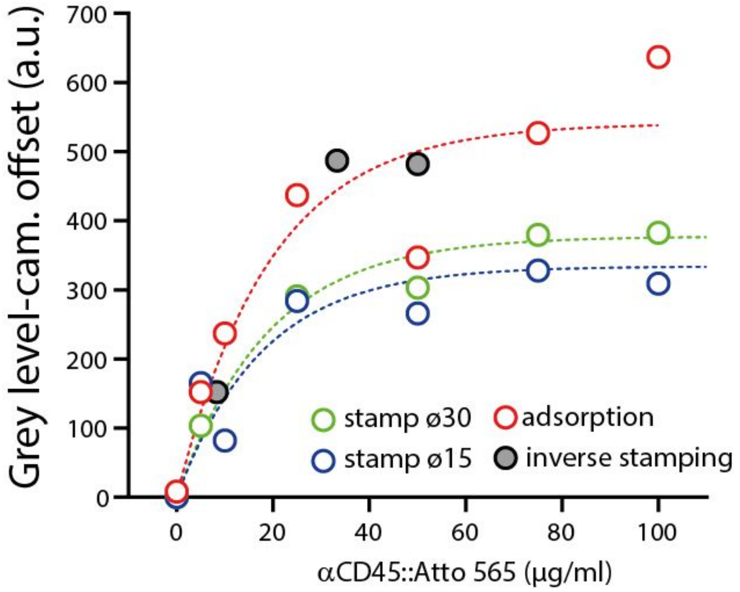
Grey level-cam. offset (a.u.)
700
600
500
400
300
200
100
0
0
20
40
60
80
100
αCD45::Atto 565 (µg/ml)
stamp ø30
adsorption
stamp ø15
inverse stamping

Figure S2

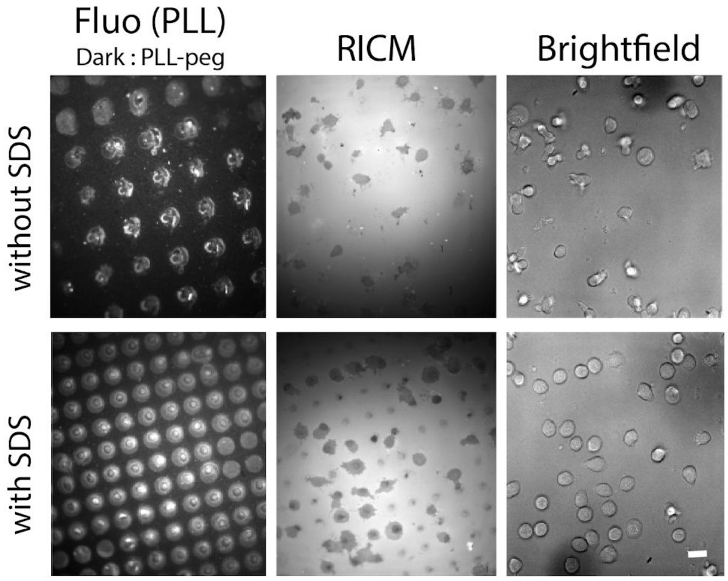
